# Deep-coverage whole genome sequences and blood lipids among 16,324 individuals

**DOI:** 10.1101/224378

**Authors:** Pradeep Natarajan, Gina M. Peloso, S. Maryam Zekavat, May Montasser, Andrea Ganna, Mark Chaffin, Amit V. Khera, Wei Zhao, Jonathan M. Bloom, Jesse M. Engreitz, Jason Ernst, Jeffrey R. O’Connell, Sanni E. Ruotsalainen, Maris Alver, Ani Manichaikul, W. Craig Johnson, James A. Perry, Timothy Poterba, Cotton Seed, Ida L. Surakka, Tonu Esko, Samuli Ripatti, Veikko Salomaa, Adolfo Correa, Ramachandran S. Vasan, Manolis Kellis, Benjamin M. Neale, Eric S. Lander, Goncalo Abecassis, Braxton Mitchell, Stephen S. Rich, James G. Wilson, L. Adrienne Cupples, Jerome I. Rotter, NHLBI TOPMed Lipids Working Group, Cristen J. Willer, Sekar Kathiresan

## Abstract

Deep-coverage whole genome sequencing at the population level is now feasible and offers potential advantages for locus discovery, particularly in the analysis rare mutations in non-coding regions. Here, we performed whole genome sequencing in 16,324 participants from four ancestries at mean depth >29X and analyzed correlations of genotypes with four quantitative traits – plasma levels of total cholesterol, low-density lipoprotein cholesterol (LDL-C), high-density lipoprotein cholesterol, and triglycerides. We conducted a discovery analysis including common or rare variants in coding as well as non-coding regions and developed a framework to interpret genome sequence for dyslipidemia risk. Common variant association yielded loci previously described with the exception of a few variants not captured earlier by arrays or imputation. In coding sequence, rare variant association yielded known Mendelian dyslipidemia genes and, in non-coding sequence, we detected no rare variant association signals after application of four approaches to aggregate variants in non-coding regions. We developed a new, genome-wide polygenic score for LDL-C and observed that a high polygenic score conferred similar effect size to a monogenic mutation (~30 mg/dl higher LDL-C for each); however, among those with extremely high LDL-C, a high polygenic score was considerably more prevalent than a monogenic mutation (23% versus 2% of participants, respectively).

---

Plasma lipids, including total cholesterol, low-density lipoprotein cholesterol (LDL-C), high-density lipoprotein cholesterol (HDL-C), and triglycerides, are heritable risk factors for atherosclerotic cardiovascular disease.^1,2^ Understanding the inherited basis for plasma lipid levels has led to new treatments and to tests to identify individuals at risk for disease. Advances in technologies to characterize DNA sequence variants (i.e., Sanger sequencing, genotyping arrays, exome sequencing) have progressively allowed us to solve monogenic forms of dyslipidemia and to uncover common DNA sequence variants as well as rare mutations that contribute to plasma lipid levels in the population. However, due to the inherent limitations of genotyping arrays and exome sequencing, the non-coding regions of the genome remains incompletely characterized, particularly for rare mutations. In addition, the relative contributions of common DNA sequence variants and rare coding mutations to extreme lipid values has not been delineated.

It is now possible to directly enumerate the whole genome sequences (WGS) of large number of individuals. When performed at sufficient depth of coverage (>20-fold coverage per base), WGS can detect single nucleotide polymorphisms, insertions, and deletions across the allele frequency spectrum in both non-coding and coding regions. These advances allow us to test the incremental value of WGS as a tool for locus discovery and also develop a framework to understand why a specific individual might have an extreme lipid value. Towards these two goals, we studied whole genome sequences in 16,324 participants of European, African, East Asian, and Hispanic ancestries with available plasma lipids phenotypes.

## Deep-coverage whole genome sequencing of 16,324 participants

Participants of the Framingham Heart Study (FHS), Old Order Amish (OOA), Jackson Heart Study (JHS), Multi-Ethnic Study of Atherosclerosis (MESA), FINRISK Study (FIN), and Estonian Biobank (EST) underwent WGS (**Fig. 1**). Following quality control (**Supplementary Table 1**), 16,324 with plasma lipids available were included in analysis (**Supplementary Table 2**). The mean (standard deviation) age was 51 (15) years and 8,669 (53%) were women. 5,911 (36%) of participants were of non-European ancestry (**Supplementary Table 2, Supplementary Fig. 1A-C**. The proportion of individuals on lipid-lowering medications was low (9%).

**Fig. 1:**
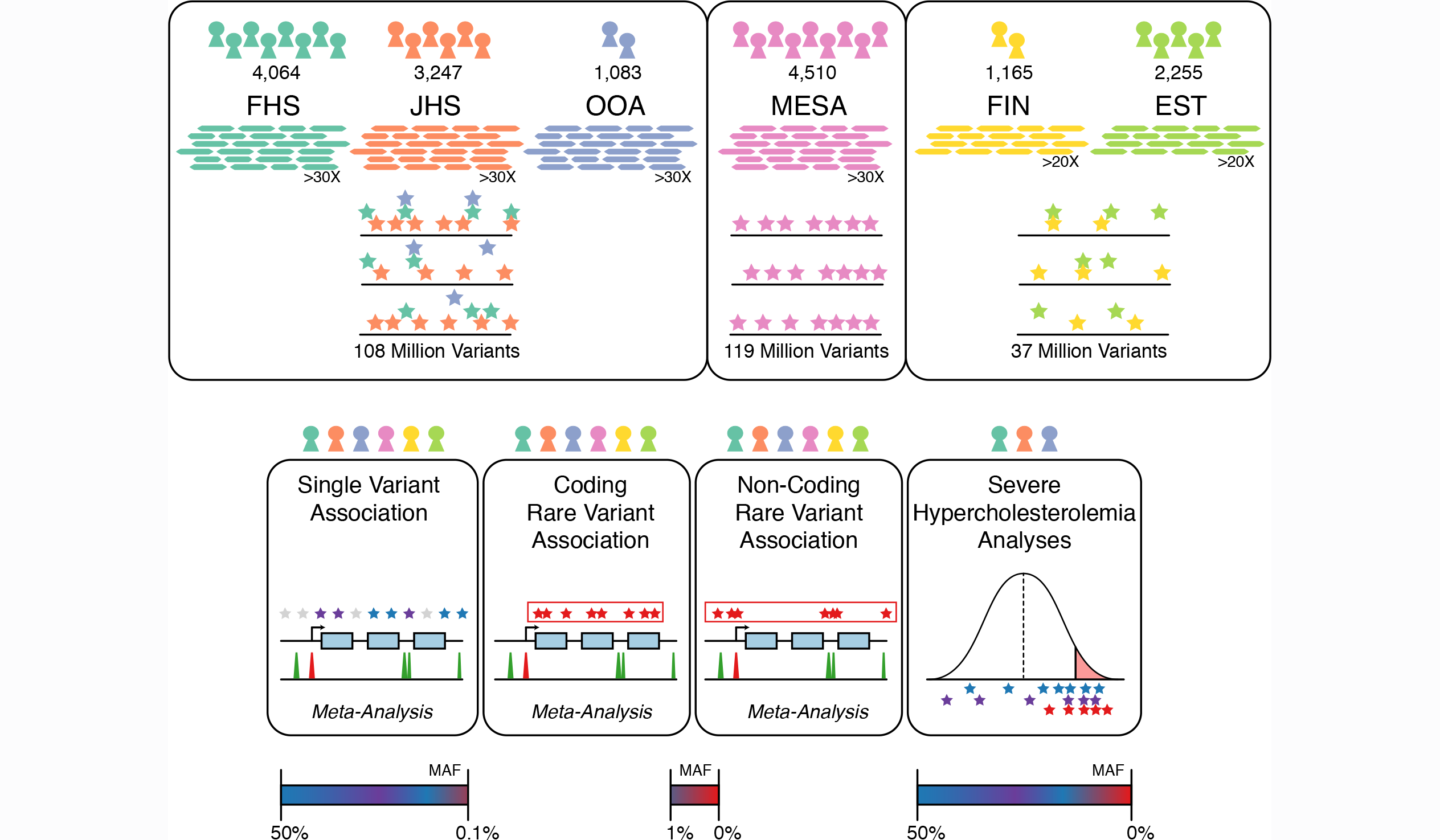
Schematic of genomic variant discovery and analyses. Variants were jointly discovered in three distinct sets: 1) FHS, JHS, and OOA; 2) MESA; and 3) EST and FIN. Cohorts included in analyses are denoted by color-coded icons. Allele frequency space assessed are indicated for analyses. EST = Estonia, FHS = Framingham Heart Study, FIN = Finland, JHS = Jackson Heart Study, MESA = Multi-Ethnic Study of Atherosclerosis, OOA = Old Order Amish

WGS target coverage was >30X for FHS, OOA, JHS, and MESA (as a part of the NIH/NHLBI Trans-Omics for Precision Medicine (TOPMed) research program) and was >20X for EST and FIN (**Supplementary Fig. 2**). The mean (standard deviation, SD) attained coverage for >30X target samples was 37.1(5.4)X and for >20X target was 29.8(5.4)X.

After performing quality control, a total of 189 million unique variants were discovered across all datasets. Total variant count characteristics varied by cohort due to sample sizes, relatedness, ethnicity, and population history (**Fig. 2**). As expected, the MESA cohort, of largely unrelated individuals of four diverse ethnicities, had the most variants per individual while the OOA cohort, a founder population of European ancestry, had the fewest variants per individual (**Supplementary Table 3**). The median number of variants per individual was 3,391,000, of which on average 4,878 were observed in only a single individual.

**Fig. 2:**
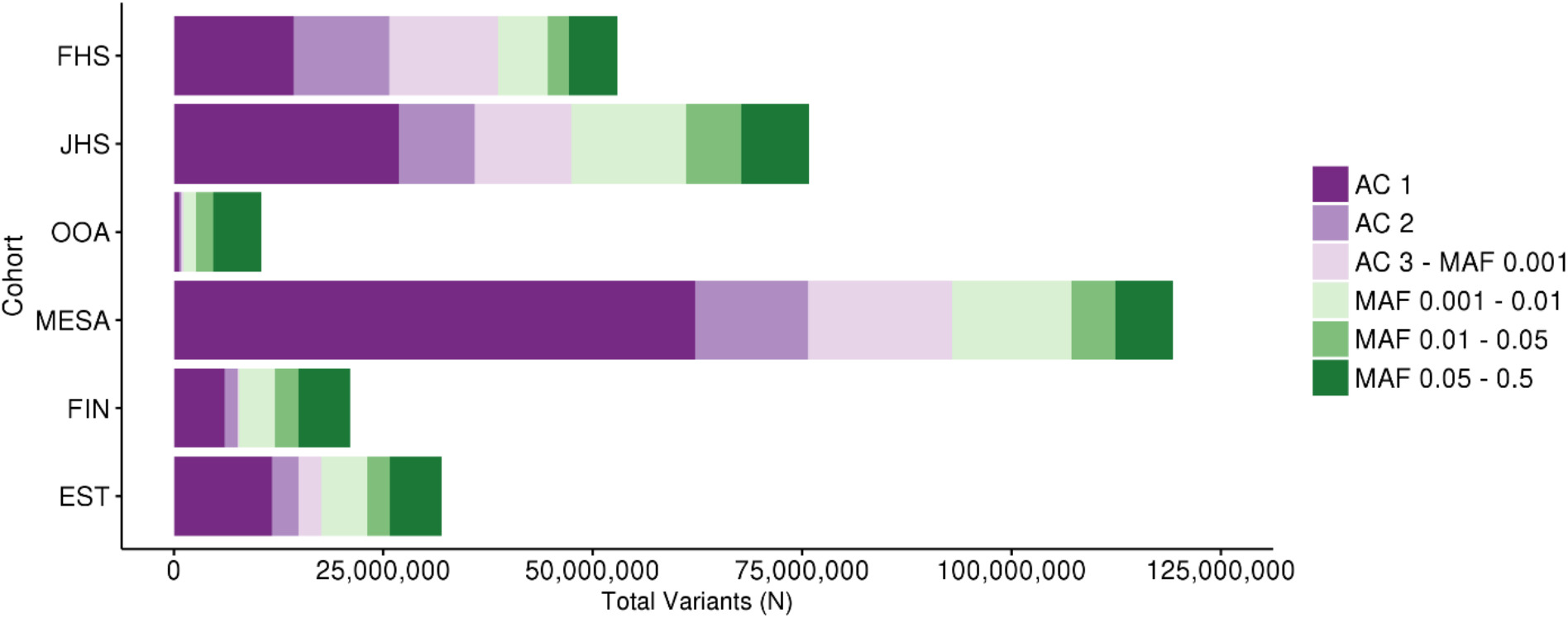
Deep-coverage whole genome sequencing identifies genomic variation across the allelic spectrum. Variant counts by allele count/frequency bin within each of the cohorts. Variants were jointly discovered in three distinct sets: 1) FHS, JHS, and OOA; 2) MESA; and 3) EST and FIN. AC = allele count, EST = Estonia, FHS = Framingham Heart Study, FIN = Finland, JHS = Jackson Heart Study, MAF = minor allele frequency, MESA = Multi-Ethnic Study of Atherosclerosis, OOA = Old Order Amish

## Common variant association study

We first analyzed common variants, i.e., those that occur often enough that it is practical to test each variant individually. We considered variants that had MAF > 0.1% within at least one of the three WGS variant callsets (minor allele count >16 for a callset that included FHS/OOA/JHS, >9 for MESA, and >6 for FIN and EST) (**Fig. 1**). We associated such variants with each of the four plasma lipids within each callset and performed inverse-variance weighted meta-analysis. Overall, 32,086,348 variants were included in this analysis. Systemic inflation was not observed (**Supplementary Table 4 and Supplementary Fig. 3A-D**). We used a conventional statistical threshold for genome-wide significance (alpha = 5 x 10^-8^)^3^ (**Supplementary Fig. 3E-H**). We observed 592, 697, 447, and 522 variants associated with total cholesterol, LDL-C, HDL-C, and triglycerides, respectively (**Supplementary Table 5**).

These variants were distributed at 10, 7, 13, and 9 loci previously associated with total cholesterol, LDL-C, HDL-C, and triglycerides, respectively, and 5 at putative novel lipid loci (**Supplementary Table 6**).^4–7^ Of the variants at known loci, 12 (38.7%) were lead variants in prior associations, 8 (25.8%) new lead variants were in high linkage disequilibrium (LD) (r^2^ > 0.8) with prior lead variants, and the remaining 11 (35.5%) new lead variants were in low LD with prior lead variants.

At a conventional alpha threshold of 5 x 10^-8^, we discovered 5 associations at putative novel lipid loci (**Supplementary Table 6**). For example, rs3215707 (MAF 2.0%), a 1-bp deletion at 9p24.1, was associated with HDL-C (+3.3 mg/dL, *P* = 1.3 x 10^-8^). rs3215707 occurs within an intron of *PLGRKT* and overlies active promoter and strong enhancer histone modification signals for hepG2 cells (**Supplementary Fig. 4**). The deletion is not in LD with SNPs and thus the association was not detectable by prior genome-wide association analyses. Within each callset, estimated effects were consistent (heterogeneity *P* = 0.53) and all demonstrated at least nominal association (P < 0.05) (**Supplementary Table 7**).

We performed iterative conditional analyses to identify distinct independent associations among 16 loci jointly reaching *P* < 5x10^-8^ for LDL-C, HDL-C, and triglycerides. While only 4 (25%) loci displayed evidence of allelic heterogeneity at *P* < 5x10^-8^, 13 (81.3%) had at least moderate evidence (P < 1x10^-4^) of allelic heterogeneity (**Supplementary Table 8**). Through conditional analyses for LDL-C, we identified a low-frequency haplotype specific to African Americans (MAF 0.1% FHS, 0% OOA, 1.0% JHS,) including variants in LD (r^2^ > 0.8) at a transcriptional transition region within the first intron of *LDLR* (rs17242843), *LDLR* promoter (rs17249141), and enhancer 4kb upstream from the *LDLR* transcription start site (rs114197570) (**Supplementary Fig. 5, Supplementary Fig. 6**). Presence of these variants resulted in a 28 mg/dL lowering of LDL cholesterol (P = 2x10^-11^), indicating a gain-of-function effect on *LDLR* (**Supplementary Fig. 7**).

## Rare variant association study of coding variants

Rare variants occur too infrequently to allow association tests of individual variants and thus, require aggregating rare variants into sets and testing quantitative trait distribution among carriers of a set versus non-carriers.^8^ We aggregated coding sequence variants within each gene that were predicted to lead to loss of function (e.g. nonsense, canonical splice-site, or frameshift) or annotated as “disruptive” by the ensemble MetaSVM^9^ *in silico* approach. The median [interquartile range] combined MAF per gene was 0.25% [0.090-0.69%] (**Supplementary Fig. 8**). To account for known bidirectional effects of disruptive mutations in some Mendelian dyslipidemia genes, we accordingly used a mixed model sequence kernel association test (SKAT).^10,11^ Five genes associated with lipids at an exome-wide level (alpha = 0.05 / ~20,000 protein-coding genes = 2.5 x 10^-6^) (*LDLR, PCSK9,* and *APOE* for LDL-C, *LCAT* for HDL-C, and *APOC3* for triglycerides). Each has been previously established as a cause of Mendelian forms of dyslipidemia (**Supplementary Table 9**).

## Rare variant association study of non-coding variants

Next, we sought to determine whether rare variants in non-coding regions associate with plasma lipids. We used four approaches to aggregate rare, non-coding variants. (**Fig. 3**). First, we aggregated variants within “sliding windows” of 3kb in length.^12,13^ Second, we connected a non-coding variant to a gene if it resided in a segment annotated as an enhancer (and within 20kb of a gene) or a region annotated as a promoter (and within 5kb of the transcription start site of a gene, TSS). Third, using gene expression information we connected a non-coding variant to a gene if it resided in a region annotated as an enhancer. Finally, we connected a non-coding variant to a gene based on a model which predicted gene-enhancer pairs using a chromatin-state model that we previously developed.^14^ Regulatory annotations were derived from the ENCODE and NIH Roadmap projects for two cell types - HepG2 and adipose nuclei - relevant to lipoprotein metabolism. For these analyses, we consider a P < 0.05 / 254,032 groups = 2.0x10^-7^ as significant (**Supplementary Table 10, Supplementary Table 11**).

**Fig. 3:**
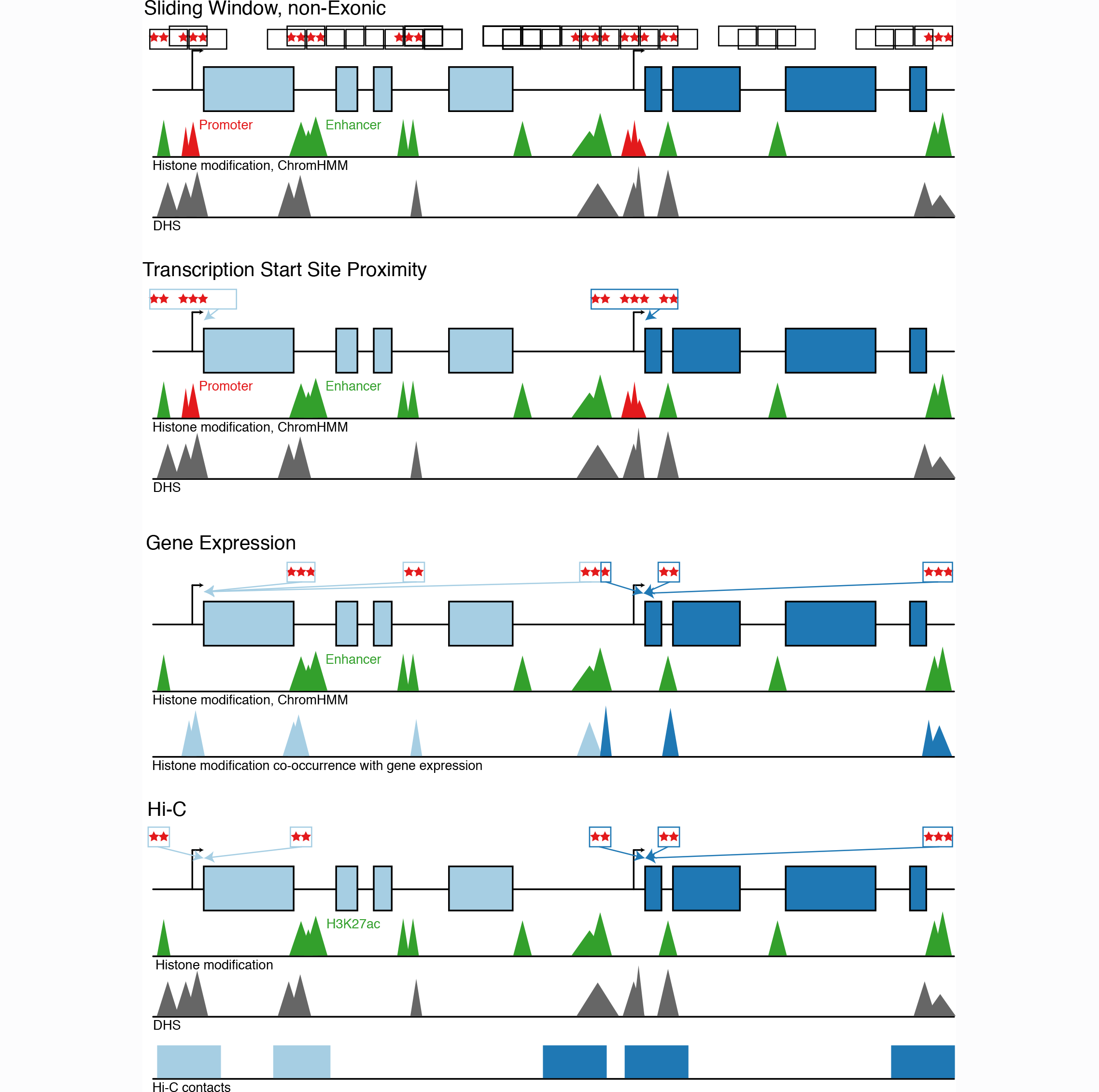
Schematic of non-coding rare variant analyses. Four grouping schematics of rare non-coding variants (MAF < 1%). The sliding window approach tiles across the genome at fixed widths, only including variants overlying annotations consistent with enhancers, promoters, and DHS in non-exonic regions. All other approaches attempt to map non-coding putative functional genomic regions with discrete genes as the analytical unit. Overall, they are based on: 1) promoter, enhancer, and DHS annotations near a gene’s transcription start site, 2) co-occurrence of enhancer and DHS annotations with HepG2 gene expression, and 3) H3K27ac marks within Hi-C contact regions mapped to genes. DHS = DNase hypersensitivity site, MAF = minor allele frequency

Using the sliding window approach, we observed suggestive associations for 3kb windows at the *CETP* (start chr16:56667000) locus (minimum *P* = 4x10^-6^) and at the *APOA1-APOC3* (start chr11:117094500) locus (minimum *P* = 8x10^-6^) with HDL-C. 17.6% of non-coding sliding windows occurring within 1Mb of known lead lipid variants were at least nominally (P < 0.05) associated with lipids versus 4.4% in other regions of the genome across all traits (P difference = 8 x 10^-272^).

An aggregation of rare non-coding variants at only two genes - *LDLR* and *APOE* - associated with LDL-C and total cholesterol (P < 2x10^-7^) (**Supplementary Fig. 9**) (**Supplementary Table 12**). The strongest *LDLR* signal (P = 9.7x10^-11^) was seen for an analysis that connected enhancers and promoters to a gene based on physical proximity (approach #2 above). Closer inspection of the specific variants shows that this signal is driven by the low-frequency haplotype specific to African Americans that was described earlier (**Supplementary Fig. 10**) (**Supplementary Table 12**). The strongest *APOE* signal (P = 8.1x10^-26^) was observed in the model connecting enhancers to a gene by gene expression (approach #3 above). Accounting for the strongest common variant association at the locus (rs7412, the *APOE e2* isoform allele), this signal attenuates to non-significance (P = 1.8x10^-2^), suggesting that the non-coding variants are driven by the *APOE e2* isoform. Beyond these two results, we found no additional signals for a burden of non-coding variants.

## Contribution of monogenic and polygenic models to extreme LDL-C values

With the availability of sequence in both coding and non-coding regions in the same samples, we estimated the simultaneous contribution of monogenic and polygenic determinants to extreme LDL-C in individuals of European (EA) and African (AA) ancestry. We defined ‘extreme’ as the top or bottom 5^th^ ancestry-specific percentile of LDL-C. Analyses were conducted in FHS and MESA-EA subjects (extremes defined as LDL-C >183 mg/dl or LDL-C <72.9 mg/dl) and JHS and MESA-AA subjects (extremes defined as LDL-C >198.6 mg/dl or LDL-C <71 mg/dl), separately.

Among participants with extremely high LDL-C, we searched for mutations in any of six Mendelian genes previously linked to high LDL-C *(LDLR, APOB, PCSK9, ABCG5, ABCG8*, and *LDLRAP1)* (**Supplementary Table 13**).

To determine polygenic contribution, we implemented a systematic approach to derive, test, and validate a new ‘genome-wide’ polygenic score for LDL-C using mutually independent datasets. A polygenic score provides a quantitative assessment of the cumulative risk associated with multiple common risk alleles for each individual. Scores for each individual participant are created by adding up the number of risk alleles at each variant and then multiplying the sum by the literature-based effect size.

We derived a genome-wide polygenic score based on the association statistics of all available common (MAF ≥ 0.01) SNPs with LDL-C, as determined by our previously published genome-wide association study.^7^ The correlation between the variants was assessed using the European reference population from the 1000 Genomes study.^17^ The LDPred computational algorithm was then used to construct genome-wide polygenic scores.^15^ This Bayesian approach calculates a posterior mean effect size for each variant based on a prior (association with LDL-C in a previously published study) and subsequent shrinkage based on the extent to which this variant is correlated with similarly associated variants in a reference population. The underlying Gaussian distribution additionally considers the fraction of causal (e.g. non-zero effect sizes) markers. Because this fraction is unknown for any given disease, LDpred uses a range of plausible values to construct different polygenic scores. We also applied various r2 and p-value thresholds to the previously published results. As a comparison to the expanded polygenic scores, we generated an additional polygenic score restricted to lead variants (P < 5x10^-8^) at distinct genomic loci, weighted by discovery beta (restricted score). The best score was determined based on maximal model fit (R2) from a linear regression models in a health-care biobank of 25,534 unrelated individuals (Nord-Trøndelag Health Study, HUNT)^16^ (**Supplementary Table 14**).

For LDL-C, a genome-wide polygenic score incorporating 2 million single nucleotide polymorphisms with LDpred provided the best model fit (**Supplementary Table 15**). We applied this polygenic score separately within the WGS samples in FHS, JHS, and MESA. We labelled individuals as high polygenic score if they fell in the top 5^th^ percentile of race-specific scores (**Table 1**).

**Table 1.** Delineating the monogenic and polygenic contributions to extremely high or low LDL cholesterol concentrations. (a). Effect of monogenic mutation or polygenic score on odds for extremely high or low LDL-C. Values are represented as OR [95% CI] for association with given trait. (b). Effect of monogenic mutation or polygenic score on LDL-C in mg/dl. Values are represented as beta [95% CI] in mg/dl for LDL-C. Multi-variable associations were performed with sex + age + age^2^ (effects not listed) with monogenic carrier status + high polygenic score using logistic (a) and linear regression (b). Polygenic risk score was derived from 2 million variants using LDpred. High polygenic score was defined as membership in the top 5^th^ percentile of the ancestry-specific score distribution. EA, European American; AA, African American.

| Extremely High LDL-C |  |  |  |  |  |  |  |  |  |  |
| --- | --- | --- | --- | --- | --- | --- | --- | --- | --- | --- |
| Ancestry | N<sub>total</sub> | N<sub>extreme</sub> | Monogenic carrier (N<sub>extreme</sub>) | High polygenic score (N<sub>extreme</sub>) | Monogenic carrier OR (95% CI) | Monogenic carrier p-value | Monogenic carrier PAF | High polygenic score OR (95% CI) | High polygenic score p-value | Top 5<sup>th</sup> percentil of Polygenic score PAF |
| EA | 5910 | 284 | 5 | 64 | 10.92 (3.71,32.14) | 1.4x10<sup>-5</sup> | 1.60 | 7.65 (5.56,10.52) | 5.7x10<sup>-36</sup> | 19.6 |
| AA | 4380 | 217 | 7 | 29 | 7.43 (3.01,18.35) | 1.4x10<sup>-5</sup> | 2.79 | 3.2 (2.1,4.89) | 6.7x10<sup>-8</sup> | 9.2 |
| Extremely Low LDL-C |  |  |  |  |  |  |  |  |  |  |
| Ancestry | N<sub>total</sub> | N<sub>extreme</sub> | Monogenic carrier (N<sub>extreme</sub>) | Low polygenic score (N<sub>extreme</sub>) | Monogenic carrier OR (95% CI) | Monogenic carrier p-value | Monogenic carrier PAF | Low polygenic score OR (95% CI) | Low polygenic score p-value | Bottom 5<sup>th</sup> percentil of Polygenic score PAF |
| EA | 5910 | 286 | 6 | 82 | 21.73 (6.2,76.15) | 1.5x10<sup>-6</sup> | 2.00 | 10.38 (7.69,14.02) | 1.5x10<sup>-52</sup> | 25.9 |
| AA | 4380 | 218 | 11 | 32 | 13.83 (6.25,30.62) | 9.4x10<sup>-11</sup> | 4.68 | 3.7 (2.46,5.58) | 3.9x10<sup>-10</sup> | 10.7 |

**Monogenic mutation or high polygenic score**
| Ancestry | N<sub>total</sub> | Monogenic carrier (N) | High Polygenic score (N) | Monogenic carrier Beta mg/dl | Monogenic carrier SE | Monogenic carrier p-value | High Polygenic score Beta mg/dl | High Polygenic score SE | High Polygenic score p-value |
| --- | --- | --- | --- | --- | --- | --- | --- | --- | --- |
| EA | 5910 | 18 | 297 | 29.98 | 8.07 | 2.1x10<sup>-4</sup> | 33.07 | 2.05 | 1.7x10<sup>-57</sup> |
| AA | 4380 | 25 | 220 | 41.05 | 7.93 | 2.3x10<sup>-7</sup> | 16.96 | 2.74 | 6.4x10<sup>-10</sup> |

**Monogenic mutation or low polygenic score**
| Ancestry | N<sub>total</sub> | Monogenic carrier (N) | Low Polygenic score (N) | Monogenic carrier Beta mg/dl | Monogenic carrier SE | Monogenic carrier p-value | Low polygenic score Beta mg/dl | Low polygenic score SE | Low polygenic score p-value |
| --- | --- | --- | --- | --- | --- | --- | --- | --- | --- |
| EA | 5910 | 12 | 297 | -47.25 | 9.55 | 7.7x10<sup>-7</sup> | -35.00 | 2.00 | 7.9x10<sup>-67</sup> |
| AA | 4380 | 28 | 220 | -41.41 | 7.47 | 3.1x10<sup>-8</sup> | -20.41 | 2.74 | 1.1x10<sup>-13</sup> |

Among EA participants, a monogenic mutation was associated with an odds ratio of 10.92 (95% CI 3.71-32.14) for extremely high LDL-C whereas a high polygenic score associated with an odds ratio of 7.65 (95% CI 5.56-10.52). In EA individuals, those who carried a monogenic mutation had 30 mg/dl higher LDL-C (when compared with non-carriers; *P* = 2.1x10^-4^) and those who had a high polygenic score had 33 mg/dl greater LDL-C (when compared with all others; *P* = 1.7x10^-57^). Of 287 EA participants with extremely high LDL-C, 2% carried a monogenic mutation and 23% had a high polygenic score.

Among AA participants, a monogenic mutation was associated with an odds ratio of 7.43 (95% CI 3.01-18.35) for extremely high LDL-C whereas a high polygenic score associated with an odds ratio of 3.2 (95% CI 2.1-4.89). In AA individuals, those who carried a monogenic mutation had 41 mg/dl higher LDL-C (when compared with non-carriers; p = 2.3x10^-7^) and those who had a high polygenic score had 17 mg/dl greater LDL-C (when compared with all others; *P* = 6.4x10^-10^). Of 217 AA participants with extremely high LDL-C, 3% carried a monogenic mutation and 13% had a high polygenic score.

We replicated the association between a high polygenic score and extremely high LDL-C in an independent sample, the ARIC cohort. Among ARIC-EA individuals, a high polygenic score was associated with an odds ratio of 7.35 (95% CI 5.95-9.10; *P* < 2x10^-16^) for extremely high LDL-C and 42.8 mg/dl (95% CI 40.0-47.5; *P* < 2x10^-16^) higher LDL-C compared with individuals without a high polygenic score. Among ARIC-AA participants, a high polygenic score was associated with an odds ratio of 2.7 (95% CI 1.77-4.09; *P* < 3.3x10^-6^) for extremely high LDL-C and a 23.2 mg/dl (95% CI 15.0-31.5; *P* = 3.8x10^-8^) higher LDL-C compared with individuals without a high polygenic score.

We analyzed the monogenic and polygenic contribution to extremely low LDL-C in EA and AA participants and found similar patterns where monogenic mutations as well as a polygenic score conferred similar effect sizes (**Table 1**).

## Discussion

Here, we performed whole genome sequencing in 16,342 ethnically diverse individuals and analyzed the incremental value of WGS for locus discovery and for clinical interpretation. We replicated associations for 28 common variant loci previously associated with lipids in much larger genome-wide association analyses. We identified an association for a low frequency 1-bp deletion at 9p24.1 with HDL-C. While we replicated burden associations of rare coding mutations at known Mendelian lipid genes, we did not detect any burden associations of rare non-coding mutations through four different approaches. Lastly, we developed a new genome-wide polygenic score and showed that such a score confers an effect size on LDL-C similar to carrying a monogenic mutation, but is also notable for a much greater frequency.

These results permit several conclusions. Using whole genome sequencing as a discovery tool, the incremental yield of new loci was modest. In particular, despite a genome-wide search using four different aggregation approaches and regulatory annotations from two relevant tissues, we identified no burden-of-rare-variant signals in non-coding regions. Mutation target size and natural selection pressure is smaller in non-coding regions when compared with coding regions; based on these considerations, power calculations have suggested that sample sizes may need to considerably larger to identify rare variant burden associations in non-coding regions.^8^

Of great interest, we observed that the relative contribution of polygenic score to extremely high LDL-C is considerably greater than monogenic mutations. For example, in EA individuals, both high polygenic score and a monogenic mutation confer similar effects (~30 mg/dl higher LDL-C) but a high polygenic score in present in 23% of participants with extremely high LDL-C whereas a monogenic mutation is present in only 2%. In most individuals who carry diagnosis of familial hypercholesterolemia, no monogenic mutation is identified with clinical exome sequencing^17,18^; for a large fraction of these ‘mutation-negative’ familial hypercholesterolemia, high polygenic scores may be operative.

Important caveats and limitations should be considered. First, appropriate definitions of statistical significance for WGS association analyses have not been harmonized in the field. The convention of alpha=5x10^-8^ comes from the assumption of performing 1,000,000 independent tests. Based on our findings and simulations from others^3^, 10^-9^ may be more appropriate for analyses across diverse ethnicities to allele frequency 0.1%. Second, power is somewhat diminished with our rare variant meta-analysis approach to combine *P* values with Fisher’s method. Given known diverse coding mutations within Mendelian genes with bidirectional effects and the inability to assume unidirectional effects within the non-coding space, we employed a SKAT statistical framework. Prior approaches leveraging covariance matrices for SKAT meta-analysis were computationally inefficient for the dataset and multiple grouping strategies.^19,20^ Thus, our approach is conservative. Third, the polygenic scores described here were derived from genome-wide association studies performed largely in EA ancestry participants.^7^ Because allele frequencies, linkage disequilibrium patterns, and effect sizes of common polymorphisms vary by ancestry, the predictive capacity of polygenic score was attenuated in non-European ancestry individuals.^21^. This is an important limitation for the field that requires efforts to characterize common genomic variation influencing complex traits among non-Europeans.

In summary, we present a large-scale WGS analysis of plasma lipids in 16,324 ethnically diverse participants. Common, non-coding variants and rare, coding variants contribute to plasma lipid variation; however, association signals for rare, non-coding mutations were not detectable.

## ACKNOWLEDGEMENTS

Whole genome sequencing (WGS) for the Trans-Omics in Precision Medicine (TOPMed) program was supported by the National Heart, Lung and Blood Institute (NHLBI). WGS for “NHLBI TOPMed: Whole Genome Sequencing and Related Phenotypes in the Framingham Heart Study” (phs000974.v1.p1) was performed at the Broad Institute of MIT and Harvard (HHSN268201500014C). WGS for “NHLBI TOPMed: The Jackson Heart Study” (phs000964.v1.p1) was performed at the University of Washington Northwest Genomics Center (HHSN268201100037C). WGS for “NHLBI TOPMed: Genetics of Cardiometabolic Health in the Amish” (phs000956.v1.p1) was performed at the Broad Institute of MIT and Harvard (3R01HL121007-01S1). WGS for “NHLBI TOPMed: Multi-Ethnic Study of Atherosclerosis (MESA)” (phs001416.v1.p1) was performed at the Broad Institute of MIT and Harvard (3U54HG003067-13S1). Centralized read mapping and genotype calling, along with variant quality metrics and filtering were provided by the TOPMed Informatics Research Center (3R01HL-117626-02S1). Phenotype harmonization, data management, sample-identity QC, and general study coordination, were provided by the TOPMed Data Coordinating Center (3R01HL-120393-02S1). We gratefully acknowledge the studies and participants who provided biological samples and data for TOPMed.

The Framingham Heart Study has been supported by contracts N01-HC-25195 and HHSN268201500001I and grant R01 HL092577. The Framingham Heart Study thanks the study participants and the multitude of investigators who over its 70 year history continue to contribute so much to further our knowledge of heart, lung, blood and sleep disorders and associated traits. The Jackson Heart Study (JHS) is supported and conducted in collaboration with Jackson State University (HHSN268201300049C and HHSN268201300050C), Tougaloo College (HHSN268201300048C), and the University of Mississippi Medical Center (HHSN268201300046C and HHSN268201300047C) contracts from the National Heart, Lung, and Blood Institute (NHLBI) and the National Institute for Minority Health and Health Disparities (NIMHD). The authors also wish to thank the staffs and participants of the JHS. Dr. Wilson is supported by U54GM115428 from the National Institute of General Medical Sciences.

This analysis was supported by the American Heart Association 17SDG33680041 (P.N.), the National, Heart, Lung, and Blood Institute of the US National Institutes of Health grants K01 HL125751 (G.M.P.), R01 HL127564 (C.W., S.K.), and TOPMed analysis support grant (G.M.P., P.N.), the Ofer and Shelly Nemirovsky Research Scholar award from Massachusetts General Hospital (S.K.), and the Donovan Family Foundation (S.K.). The sponsors had no role in the design and conduct of the study; collection, management, analysis, and interpretation of the data; preparation, review, or approval of the manuscript; and decision to submit the manuscript for publication.

## AUTHOR CONTRIBUTIONS

Concept and design: P.N., G.M.P., S.M.Z., G.A., J.G.W., L.A.C., C.J.W., S.K., Acquisition, analysis, or interpretation of data: P.N., G.M.P., S.M.Z., M.M, A.G., M.C., A.V.K., W.Z., J.B., J.R.O., S.E.R., M.A., J.A.P., I.L.S., T.E., S.R., A.C., B.N., G.A., B.M., S.S.R., J.G.W., L.A.C., J.I.R., C.J.W., S.K., Drafting of the manuscript: P.N., G.M.P., S.M.Z., M.M., A.G., S.R., B.M., S.S.R, J.G.W., L.A.C., J.I.R., C.J.W., S.K. Administrative, technical, or material support: P.N., G.M.P., S.K.

## METHODS

### Study participants

Please refer to **Supplementary Text** for study participant details.

### Whole genome sequencing, variant calling, and genotyping

Sequencing was performed at one of four sequencing centers, with all members within a cohort sequenced at the same center. For the TOPMED phase 1 data, 4,148 FHS individuals and 1,095 OOA individuals were sequenced at the Broad Institute of Harvard and MIT (Cambridge, MA), while 3,266 JHS individuals were sequenced at University of Washington Northwest Genomics Center (Seattle, WA). 4,601 MESA individuals were additionally sequenced at the Broad Institute of Harvard and MIT as part of TOPMED Phase 2. 1,180 Finnish FINRISK individuals and 2,281 Estonian Biobank participants were sequenced at the Broad Institute of Harvard and MIT (Cambridge, MA).

TOPMED phase 1 BAM files provided by the sequencing centers were harmonized by the TOPMed Informatics Research Center (IRC) before joint variant discovery and genotype calling across studies. In brief, sequence data were received from each sequencing center in the form of bam files mapped to the 1000 Genomes hs37d5 build 37 decoy reference sequence. Processing was coordinated and managed by the ‘GotCloud’ processing pipeline.^22^

The two sequence quality criteria were used in order to pass sequence data on for joint variant discovery and genotyping are: estimated DNA sample contamination below 3%, and fraction of the genome covered at least 10x 95% or above. DNA sample contamination was estimated from the sequencing center read mapping using software verifyBamId.^23^

The genotype call sets used for analysis are from “freeze 3a” of the variant calling pipeline performed by the TOPMed Informatics Research Center (Center for Statistical Genetics, University of Michigan, Hyun Min Kang, Tom Blackwell and Goncalo Abecasis). The software tools used in this version of the pipeline are available in the following repository: https://github.com/statgen/topmed_freeze3_calling. Variant detection from each sequenced (and aligned) genome is performed by vt discover2 software tool.^24^ The variant calling software tools are under active development; updated versions can be accessed at http://github.com/atks/vt or http://github.com/hyunminkang/apigenome.

WGS for MESA, FINRISK, and the Estonian Biobank was performed using the Illumina HiSeqX platform at the Broad Institute of Harvard and MIT (Cambridge, MA). DNA samples are informatically received into the Genomics Platform’s Laboratory Information Management System via a scan of the tube barcodes using a Biosero flatbed scanner. All samples are then weighed on a BioMicro Lab’s XL20 to determine the volume of DNA present in sample tubes. Following this the samples are quantified in a process that uses PICO-green flourescent dye. Once volumes and concentrations are determined the samples are then handed off to the Sample Retrieval and Storage Team for storage in a locked and monitored −20C° walk-in freezer.

Libraries were constructed and sequenced on the Illumina HiSeqX with the use of 151-bp paired-end reads for WGS and output was processed by Picard to generate aligned BAM files (to hg19).^25,26^ Samples were tracked by automated LIMS messaging. Samples were fragmented with acoustic shearing and libraries were prepared with a KAPA Biosystems kit. Libraries were normalized to 1.7 nM. Cluster amplication was performed using Illumina cBot and flowcells were sequenced in HiSeq X. Variants were discovered using the Geome Analysis Tookit (GATK) v3 HaplotypeCaller according to Best Practices.^27^ Variants from MESA samples were were generated in one callset. Finland and Estonia samples were jointly called in a separate callset.

### Whole genome sequence quality control

The following three approaches were used by the TOPMed Genetic Analysis Center to identify and resolve sample identity issues: (1) concordance between annotated sex and biological sex inferred from the WGS data, (2) concordance between prior SNP array genotypes and WGS-derived genotypes, and (3) comparisons of observed and expected relatedness from pedigrees.

The variant filtering in TOPMed Freeze 3 were performed by (1) first calculating Mendelian consistency scores using known familial relatedness and duplicates, and (2) training SVM classifier between the known variant sites (positive labels) and the Mendelian inconsistent variants (negative labels). Two additional hard filters were applied: (1) Excess heterozygosity filter (EXHET), if the Hardy-Weinberg disequilbrium p-value was less than 1x10^-6^ in the direction of excess heterozygosity. An additional ~3,900 variants were filtered out by this filter, and (2) Mendelian discordance filter (DISC), with 3 or more Mendelian inconsistencies or duplicate discordances observed from the samples. An additional ˜370,000 variants were filtered out by this filter. Functional annotation for each variant was provided in the INFO field using snpEff 4.1 with a GRCh37.75 database.^28^ Analysis used hard-call genotypes, without genotype likelihoods. Genotypes with a depth < 10 were excluded.

Additional measures for quality control of TOPMed Phase I Freeze 3 and quality control for MESA, Finland, and Estonia were performed using the Hail software package.^29^ Samples were filtered by contamination (>3.0% for all, except >5.0% for Finland and Estonia), chimeras >5%, GC dropout >4, raw coverage (<30X for all, except <19X for Finland and Estonia), indeterminant genotypic sex or genotypic/phenotypic sex mismatch.

Variants for MESA, Finland, and Estonia were initially filtered by GATK Variant Quality Score Recalibration. Additionally, genotypes with GQ<20, DP<10 or >200, and poor allele balance (homozygous with <0.90 supportive reads or heterozygous with <0.20 supportive reads) were removed. And variants within low complexity regions were removed across all samples.^30^ Variants with >5% missing calls, quality by depth <2 (SNPs) or <3 (indels), InbreedingCoeff <-0.3, and pHWE <1x10^-9^ (within each cohort) were filtered out.

### Annotation

Variants were annotated with Hail using annotations from Ensembl’s Variant Effect Predictor^31^ for protein-coding annotations and Reg2Map HoneyBadger2-intersect for regulatory annotations at DNase I regions −log_10_(*P*) >= 10 (https://personal.broadinstitute.org/meuleman/reg2map/HoneyBadger2-intersectrelease/).

### Traits

Conventionally measured plasma lipids, including total cholesterol, LDL-C, HDL-C, and triglycerides, were included for analysis. LDL-C was either calculated by the Friedwald equation when triglycerides were <400 mg/dl or directly measured. Given the average effect of statins, when statins were present, total cholesterol was adjusted by dividing by 0.8 and LDL-C by dividing by 0.7, as previously done.^32^ Triglycerides were natural log transformed for analysis.

### Single variant association analysis

Single variant analysis was performed in EPACTS with Efficient Mixed-Model Association eXpedited (EMMAX) for associating each variant site with each lipid outcome within each jointly called variant call file (VCF).^10^ The results were meta-analyzed for each trait using the METAL software. All analyses were adjusted for age, age^2^, sex, cohort, and an empirically derived kinship matrix to account for both familial and more distant relatedness. For the TOPMed Phase I VCF, which included OOA, LDL-C and total cholesterol analyses were also adjusted for *APOB* p.R3527Q and triglycerides and HDL-C analyses were also adjusted for *APOC3* p.R19Ter. To ensure robust results, we only performed single variant analysis for variants with a MAF > 0.1%. Variants were meta-analyzed across all three VCFs using METAL.^33^ Summary statistics only for variants with MAF > 0.1% for the given VCF were included in the meta-analysis. Statistical significance alpha of 5x10^-8^ was used for these analyses.

For loci with at least one variant with *P* <5x10^-8^ within the TOPMed Phase I VCF, iterative conditional association analysis was performed. Iterative conditioning was performed until *P*>1x10^-4^ was attained.

### Rare variant association analyses

#### Coding

We first identified rare (MAF <1%) mutations for each VCF within coding sequences. After Variant Effect Predictor^31^ annotation, we identified loss-of-function (e.g. nonsense, canonical splice-site, and frameshift) and disruptive missense (by MetaSVM^9^).

#### Sliding window

We also performed rare variant association tests within the non-coding space (**Supplementary Figure 7**). As before, we performed a “sliding window” approach aggregating 3kb (overlapping by 1.5kb) windows and considering rare variants occurring within enhancer or promoter elements at DNase I hypersensitivity sites.

#### Proximity to transcription start sites

For non-coding tests, we next attempted to link rare non-coding variants with genes for association testing using regulatory annotations for HepG2 and adipose nuclei from ENCODE and NIH Roadmap. Given prior observations showing enrichment of functional promoter variants at *LIPG* with HDL-C extremes^34^, we similarly aggregated variants near transcription start sites (TSS). Prior studies have shown that approximately 80% of cis-eQTLs fall within 100kb of TSS.^35^ To increase likelihood that of mapping regulatory variants to the nearest gene, we were more restrictive and included variants overlapping promoter sequences +/-5kb and enhancer sequences +/-20kb of TSS at DNase I hypersensitivity sites.

#### Linking enhancers to genes by gene expression

We also linked chromatin state defined enhancers with genes using data from the Roadmap Epigenomics project^36^ and the method presented previously^37^ with a few small modifications.^38^ The method predicts links using chromatin state information, position of the enhancer relative to the TSS, and the correlation of multiple chromatin marks with gene expression across cell types. Here we used the correlation with gene expression of the signal of five chromatin marks: H3K27ac, H3K9ac, H3K4me1, H3K4me2, and DNaseI hypersensitivity. The gene expression data was the RPKM expression data for protein coding exons across 56 reference epigenomes from the Roadmap Epigenomics project (available in the file 57epigenomes.RPKM.pc from http://compbio.mit.edu/roadmap; Universal Human Reference was excluded). The chromatin mark signal was the -log_10_(*P*) tracks averaged to a 200-bp resolution. As input to our code we used the version of those tracks first averaged at 25-bp resolution using the ‘Convert’ command of ChromImpute.^39^ In computing correlation between a specific chromatin mark signal and gene expression we used the Pearson correlation and omitted from the calculation samples lacking both chromatin mark signal and gene expression data. We made predictions separately for each of the 127 reference epigenomes and locations assigned to chromatin states, 6_EnhG, 7_Enh, and 12_EnhBiv, of the 15-state core 5-marks ChromHMM model.^36,40^ We restricted our predictions to chromatin state assignments on chr1-22 and chrX. We considered linking 200-bp bins within 1MB of a TSS of each gene as annotated in the file Ensembl_v65.Gencode_v10.ENSG.gene_info available from http://compbio.mit.edu/roadmap.^36^ If a gene had multiple TSS, then we only used the outermost TSS.

The method for linking is based on determining for each combination of cell type, chromatin state, and position relative to the TSS the estimated probability the set of correlations we observed would come from the actual data compared to randomized data. To this end we created a training set of actual observed correlations (positive examples) and correlations computed after randomizing which gene expression values were assigned to which genes (negative examples) separately for each combination of cell type, chromatin state, and position relative to the TSS. Each entry in the training set has five features corresponding to correlations for each of the considered chromatin marks. There is a positive and a corresponding negative entry for each instance of the specified chromatin state in the specified cell type at the specified position relative to the TSS or within 5kb of it (for smoothing purposes). We trained a logistic regression classifier to discriminate actual correlations with randomized correlations. We used the logistic regression library implemented in the Weka package version 3.7.3 with the regularization parameter set to 1.^41^ For considering linking a specific instance of a chromatin state assignment in a specific cell type and position relative to the TSS of a gene we applied the corresponding classifier. Let p denote the probability the classifier gives of being in the positive class of the actual observed correlations. We retained those links for which p/(1-p) was greater than or equal to 2.5. The method we used here is implemented in the code LinkingRM.java. For the analyses presented here we used those links for the primary enhancer state, 7_Enh.

#### HiC/enhancer links to genes

To connect non-coding variants with putative target genes, we predicted functional gene-enhancer pairs using a chromatin state-based model we previously developed.^14^ This model assumes that the impact of an enhancer on gene expression is determined by the product of its intrinsic “Activity” (for which we use quantitative DNase-Seq and H3K27ac ChIP-Seq levels as a proxy) and the “Contact Frequency” at which the enhancer physically encounters its target promoter in the nucleus (for which we use Hi-C data as a proxy). We previously found such an Activity by Contact (ABC) model accurately identifies enhancers whose perturbation leads to changes in gene expression in the human MYC locus^14^, and we have since found that the same model can identify enhancers across other gene loci and cell types (C. Fulco, E. Lander, and J. Engreitz, *in preparation*). We extended our previously published model to predict enhancer-gene connections in the liver, using DNase-Seq and H3K27ac ChIP-Seq data from a hepatocarcinoma cell line (HepG2) previously generated by the ENCODE project.^42^ To define putative regulatory elements, we expanded DNase-Seq peak calls from ENCODE by 500 bp on either side and merged overlapping peaks.^14^ For each element, we calculated Activity as a function of the normalized read count of H3K27ac and DNase-Seq. Because high-resolution Hi-C data is not available for HepG2 cells, we estimated the Contact probability between putative regulatory elements and genes using the average profile across deeply sequenced Hi-C libraries from 7 different cell types^43^ as previously described^14^. For each putative enhancer-gene pair, we calculated an “ABC score” equal to the Activity × Contact of the putative enhancer normalized by the sum of Activity × Contact across all other putative elements within 5 Mb of the target gene. We tuned free parameters in this model (such as the relative weight of DNase-Seq and H3K27ac data and a pseudocount to add to Hi-C data) and chose a threshold cutoff using a set of experimentally measured enhancer-promoter connections in two cell types (C. Fulco, E. Lander, and J. Engreitz, *in preparation).* This analysis defined, for each expressed gene, a set of elements predicted to regulate that gene in HepG2 cells. These sets of elements were used for gene-level variant burden tests.

#### Statistical test

We tested the association of the aggregate of the aforementioned groupings with each lipid trait using the mixed-model Sequence Kernal Association Test (SKAT) implementation in EPACTS to account for bidirectional effects.^10^ Analyses were adjusted for age, age^2^, sex, cohort, and empiric kinship. *P* values were meta-analyzed using Fisher’s method. Statistical significance for each gene-based test was 0.05 / 20,000 tests = 2.5x10^-6^.

### Lipid extremes analysis

#### Traits

We first defined LDL-C extremes as the top and bottom ancestry-specific 5^th^ percentiles from the data (LDL-C > 183 mg/dl or > 198.6 mg/dl for EA and AA, respectively; LDL-C < 72.9 mg/dl or <71 mg/dl for EA and AA, respectively).

#### Monogenic determinants

We catalogued mutations in Mendelian genes previously linked to extreme LDL-C (**Supplementary Table 13**). We included variants that were previously linked to Mendelian dyslipidemia in ClinVar (“pathogenic” or “likely pathogenic” with no “benign”) or loss-of-function, and had an allele frequency <1% (autosomal dominant) or <10% (autosomal recessive). Genotypes were only considered based on expected inheritance pattern (autosomal dominant or autosomal recessive).

#### Polygenic score derivation

We evaluated three distinct approaches to generate weighted polygenic scores using prior genome-wide association analysis summary statistics^7^: 1) only lead variants at genome-wide significant loci, 2) varying *P* and linkage disequilibrium r^2^ thresholds (defined by 1000G CEU) using PLINK^44^, and 3) all variants but adjusting weights according to *P* and r^2^ (by 1000G CEU) with LDpred varying *rho*^15^. To minimize errors from strand flips, A/T and C/G SNPs were excluded. The scores were calculated as additive sums of risk allele counts for included SNPs multiplied by weights (discovery effect estimates for 1) and 2), or adjusted by LDpred for 3)).

Polygenic scores were generated within the HUNT cohort, the training set.^16^ Lipid values were extracted from the electronic health record; absence of lipid-lowering therapy was prioritized. For each trait, the model with the best fit, as measured by R^2^, was chosen to applying to the testing set, TOPMed samples.

#### Statistical analysis

In a multivariable model, we associated likelihood of membership within the extreme tail of a trait with monogenic mutation carrier status, high (top 5^th^ percentile) or low (bottom 5^th^ percentile) polygenic score, age, age^2^, and sex, separately in European American (EA from FHS and MESA-EA) and African American (AA from JHS and MESA-AA) samples. We also ran linear regression models with continuous LDL-C and the independent variables listed above.

